# Formation process of the twinning β-form anhydrous guanine platelets in the scallop eyes

**DOI:** 10.1101/2022.11.14.516375

**Authors:** Dongmei Guo, Yiqun Liu, Xiubin Hou, Xubo Wang, Chenge Fan, Lixia Bao, Xinpeng He, Hongmei Zhang, Yurong Ma

## Abstract

Square shaped twinning guanine microplates with high symmetry are assembled into highly ordered layered patterns and function as image-forming mirrors in the scallop eyes. However, the formation process and biomineralization mechanism of twinning guanine microplatelets are still unclear. Herein, the eyes of juvenile *Yesso* scallops were investigated to understand the formation mechanism of the twinning β-form anhydrous guanine (β-AG) microplatelets exposing (100) plane. We find α form anhydrous guanine (α-AG) and single-crystal β-AG nanoplatelets in the very early stage of the eyes of the juvenile scallops, while the α-AG was supposed to be formed via amorphous guanine during the sample preparation process. Besides β-AG and α-AG, amorphous guanine was found in the eyes of juvenile scallops with size of 2.5 mm according to the Raman spectra. A formation mechanism was proposed for the biogenic twinning guanine platelets. Firstly, amorphous guanine is formed as an intermediate phase, which transforms into single crystalline β-AG nanoplatelets, or, dissolve and recrystallize to single crystalline β-AG nanoplatelets. Then, a second layer of β-AG forms on the top of the original single crystalline β-AG nanoplatelets, forming twinning β-AG nanoplatelets with two *c* axes with a certain angle, 83° or 14°. Each layer of the β-AG nanoplatelets is calculated to be about 14 ± 2 nm. This is the first time to report the formation mechanism of biogenic twinning β-AG microplatelets. Uncovering the formation mechanism of twinning platelets of organic crystals may shed light on the formation of functional synthetic twinning organic crystals in the laboratories.

## 1. Introduction

Biological organic and inorganic minerals formed in organisms usually have excellent optical,^[1]^ magnetic^[2]^ and mechanical^[3]^ properties, attributing to the elaborate hierarchically ordered micro- and nanostructures of the biominerals. Biogenic guanine crystals formed in organisms exhibit excellent optical properties such as information transmission,^[4, 5]^ structural color,^[1, 6, 7]^ broad-band and narrow-band reflectors,^[8-10]^ and vision enhancement,^[11-14]^ due to the extremely high refractive index of β-form anhydrous guanine (1.85) and exquisite control of the polymorphs, morphologies, sizes, exposed planes and highly ordered assemblies.^[9, 15, 16]^ For example, the guanine crystals in Neon Tetra,^[7]^ sapphirinid copepods,^[5]^ and chameleon^[17]^ could change the color of body surface, allowing them effectively to camouflage in changing environments. Moreover, twinning β-AG microplatelets are the important components of mirror-imaged eyes of scallops.^[18]^ The micro-nano structures and reflection properties of twinning guanine microplatelets in the concave mirror of scallop eyes have been clearly studied.^[12, 18]^ Recently, Wagner and collaborators proposed that guanine nucleates from a disordered precursor phase and the crystals grow by oriented attachment of partially ordered nanogranules into twinning platelets and then single crystals, a non-classical crystallization process.^[19]^ However, the biomineralization mechanism of the biogenic twinning guanine microplatelets is still a mystery.

Guanine has four polymorphs: α and β form anhydrous guanine with N_7_ protonated purine ring,^[20]^ as well as guanine monohydrate (GM) and dehydrated-GM with N_9_ protonated purine ring.^[21-23]^ The biogenic guanine crystals in fish scales, copepod cuticles, spider epidermis, scallop eyes and other organisms are mainly β-form anhydrous guanine (β-AG),^[8, 18, 24-26]^ while GM,^[27]^ dehydrated-GM^[22]^ and α-form anhydrous guanine (α-AG) ^[28]^ can only be synthesized in the laboratory.

The anhydrous twinning guanine platelets and low-refractive-index cytoplasmic layers are alternately arranged and form the concave mirrors for imaging in the eyes of adult *Yesso* scallops. However, the concave mirror consisted of guanine crystals is absent in the eye of very early stage of larve scallops.^[29]^ During the early stage of the scallop eye development, the concave mirror composed of β-AG microplatelets is formed between the retina and the pigment layer.^[12, 29]^ Thus, taking juvenile scallops as specimens, it is possible to discover the formation mechanism of biological twinning guanine platelets. Herein, the eyes of juvenile scallops were used as the experimental samples to explore the formation process of biological twinning guanine platelets by using Raman microscopy, TEM and SAED. We find that in the early stage of eye development, there were amorphous guanine and guanine single-crystal nanoplatelets in the region where the concave mirror would be formed. The crystallization mechanism of biological twinning guanine platelets was proposed for the first time according to the crystalline state of guanine in the scallop eyes.

## 2. Results

The guanine regions in the eyes of scallops at different development stages were characterized in detail to investigate the crystallization process of twinning guanine microplatelets.

### 2.1. Characterization of the eyes of larvae scallop (1.5 mm)

We tracked the developing process of larvae *Yesso* scallop under optical microscope and found that the eyes can be visible when the size of the scallops are larger than 1.5 mm **(Figure S1)**. As shown in Figure S1A, seven tentacles and two eyes marked with red circles can be observed on the edge of the mantle of a juvenile scallop with size of 1.5 mm. The semi-thin sections of the eye with thickness about 200 nm dissected from a 1.5 mm larvae scallop were characterized by TEM. At this stage, the eyes just start to be developed and much of the tissue such as retina, lens and concave mirror can’t be observed yet from the TEM image. The oval band marked by the dotted red line in **Figure 1**A was the region where the concave mirror composed of guanine crystals would form at a later stage **(Figure 2)**. The retina began to differentiate inside the oval band of the eye, which were marked with green arrows (Figure 1 B and C). There was a large population of undifferentiated cells in the middle of the eye. There were many short bands packed into layers on the oval band, (Figure 1D), which is probably the location where the guanine crystals would form in the later stage. The short layered bands within the oval region exhibit amorphous feature according to the selected area electron diffraction (SAED) (inset of Figure 1D).

**Figure 1.**
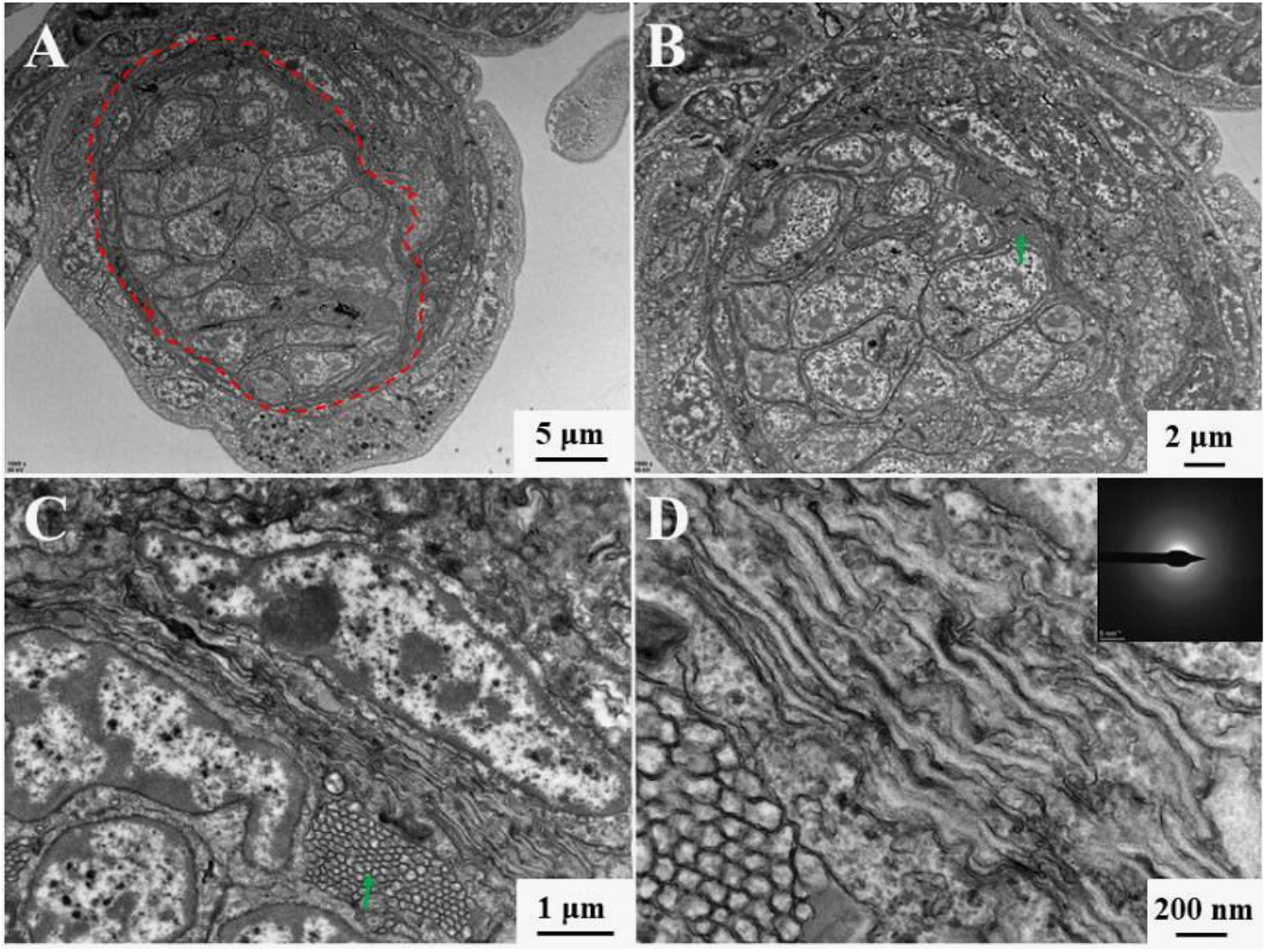
TEM images of the eye of a 1.5 mm larvae scallop. (A) The eye with low magnification, (B-D) zoomed-in images. Inset: SAED patterns of the layered short bands within the Figure (D). The red dotted line indicated the region where the concave mirror composed of guanine would be formed. The green arrow pointed to the retina.

**Figure 2.**
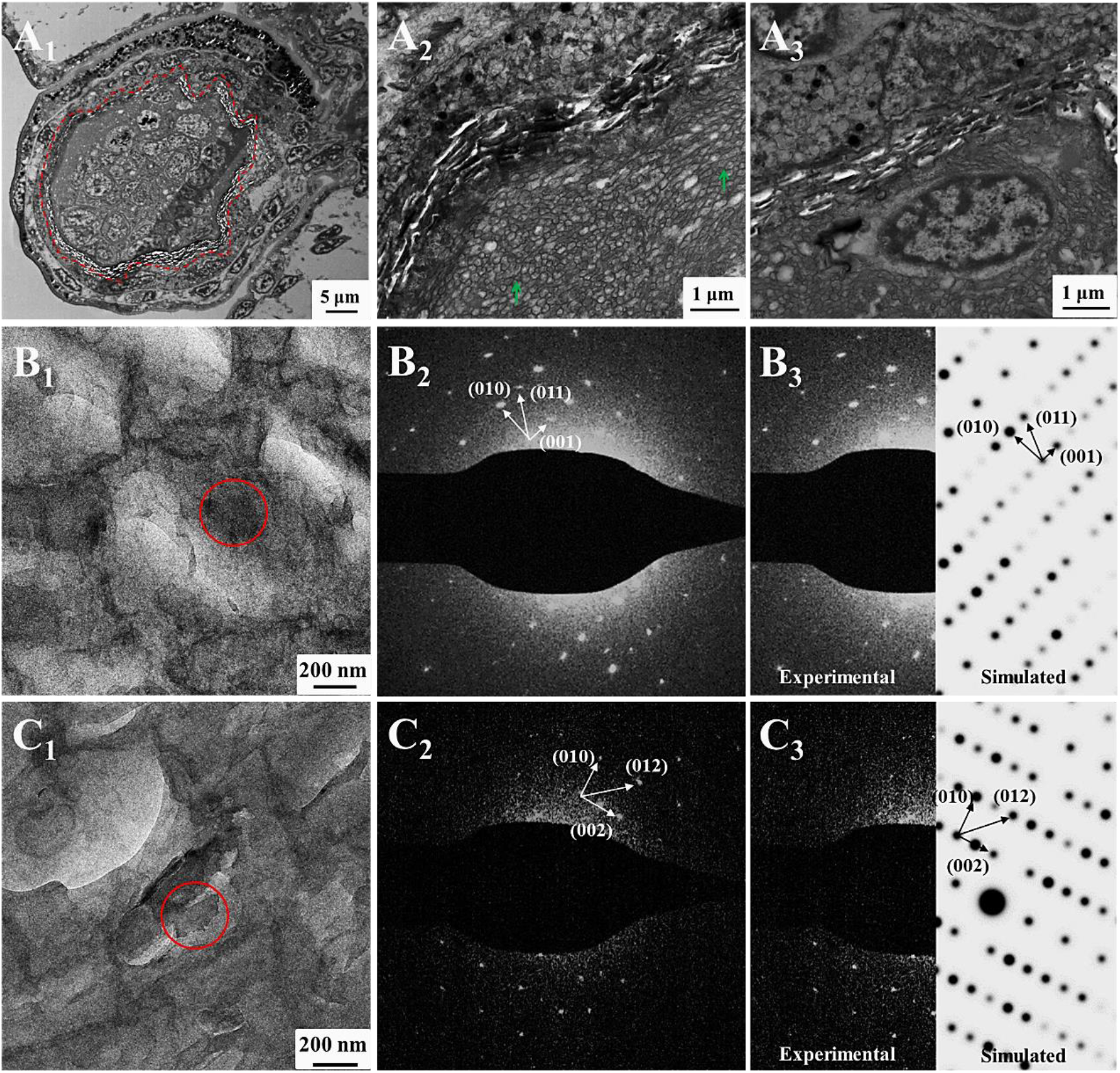
TEM images and SAED patterns of the guanine crystals in the eye of 2.5 mm juvenile scallops; the experimental SAED patterns (B_2_, C_2_) and simulated SAED patterns (B_3_, C_3_) of the guanine crystals in (B_1_, C_1_). (A_1_) The eye with full outline. (A_2_, A_3_) Zoomed-in images of the region of concave mirror in (A_1_). The red dotted line indicates the concave mirror. Most of the crystal plates on the section expose side faces, and only a few platelets exposed large crystal faces. The green arrows show the retina.

Micro-Raman spectra on the above oval band region were collected in order to further understand whether there exists guanine in the oval region of the eyes dissected from 1.5 mm larvae scallop. No any typical peak of α-AG or β-AG was detected according to the micro-Raman spectra obtained on the tested sites **(Figure S2)**. Therefore, we can conclude that no peaks of amorphous and crystalline guanine can be detected in the oval regions of the eyes of larvae scallop with size of 1.5 mm.

### 2.2. Micro-Raman spectroscopy and TEM of the eyes of juvenile scallops (2.5 mm)

The shells become opaque and the number of eyes increased significantly when the body length of the larvae scallops grows to be about 2.5 mm (Figure S1B). There were white bright spots appeared at the center of the eyes which may be caused by the light reflection of the concave mirror. The semi-thin sections of the 2.5 mm scallop eyes were also characterized by TEM and SAED. It is worthy to notify that dark rod-like materials packed as layers can be observed on the oval band area of the eyes (Figure 2 A_1_-A_3_), consistent with that concave mirror region of the adult scallops reported in the literature.^[12]^ These packed rod-like particles are supposed to be the guanine platelets in the concave mirror region. This oval band area of the eyes observed in Figure 2A_1_ is at the same position as the oval band of the eyes of larvae scallop with size about 1.5 mm (Figure 1). This result also indirectly confirms that the oval band region of the eye of mm scallop was the concave mirror region of the eyes where crystalline guanine would form in later stage of scallops. The white bands in the concave mirror region are actually holes, leftover of the loss of guanine crystals during the ultrathin microtome process. During the slicing process of ultrathin microtome, guanine crystals with high hardness may tend to be split off from the surrounding organism. Most of the crystal platelets exposed the side faces, and only a few platelets exposed a large crystal face. The retina was clearly differentiated in the inner side of concave mirror and it distributed in a much larger area than that of the 1.5mm scallops (Figure 2A_2_).

SAED of the nanoplatelets exposing large crystal face shows d-spacings 1.83 nm, 0.97 nm, 0.88 nm, corresponding to (001), (010), (011) planes of β-AG, respectively (Figure 2B). The d spacings of 0.92 nm, 0.98 nm, 0.64 nm in Figure 2C were indexed to the (002), (010), (012) faces of β-AG. The SAED patterns indicate that the guanine platelets in the eyes of 2.5mm juvenile scallop were single crystals exposing (100) plane (Figure 2B_2_, B_3_, C_2_, C_3_), and the SAED pattern is consistent with the simulated SAED pattern of β-AG with [100] zone axis. These results suggest that the zone axis of these guanine crystals was [100] of the β-AG. The (100) face was the largest crystal face exposed by the guanine crystals, and the (001) and (010) planes were in the diagonal direction of guanine single crystalline platelet, respectively. In the concave mirror region, some guanine single crystals were square, while others were rectangular. The guanine single crystals with a rectangular shape may be formed by the incomplete growth of guanine. In order to investigate the formation process and determine the crystal form of guanine crystals in the concave mirror region of scallop eyes, Raman spectra of the concave mirror region of the eyes of scallops with different growth stages were collected by using confocal Raman microscopy. The Raman spectra of pure α-AG, β-AG and synthetic amorphous guanine (AmG) synthesized in the laboratory were applied as reference spectra. The α-AG sample was synthesized in formamide with guanosine as additive, whereas the β-AG sample was obtained in formamide with hypoxanthine as additive^[28]^. Amorphous guanine was synthesized according to our previously reported rapid neutralization method.^[30]^ The characteristic peaks at 40 cm^-1^, 63 cm^-1^, 73 cm^-1^, 94 cm^-1^ and 107 cm^-1^ were assigned to α-AG, while the peaks at 39 cm^-1^, 72 cm^-1^, 107 cm^-1^ and 204 cm^-1^ were attributed to β-AG **(Figure 3**B). The synthetic amorphous guanine cannot be distinguished individually since the characteristic peaks of synthetic amorphous guanine were at 72 cm^-1^ and 107 cm^-1^, very similar to that of β-AG. Similar vibration bands of synthetic amorphous guanine and β-AG in their Raman spectra might attribute to their similar short-range ordered structures.^[30]^ All these Raman spectra of guanine samples have characteristic vibration bands of guanine at 650 cm^-1^, 940 cm^-1^ and 1230 cm^-1^ (Figure 3C), consistent with that reported in the literature^[16]^. Interestingly, the obtained Raman spectra of different positions of the concave mirror in the eye of juvenile scallops with size of 2.5 mm have the above three typical vibration bands at high wavenumbers between 650 cm^-1^ and 1300 cm^-1^, indicating the presence of guanine at these locations.

**Figure 3.**
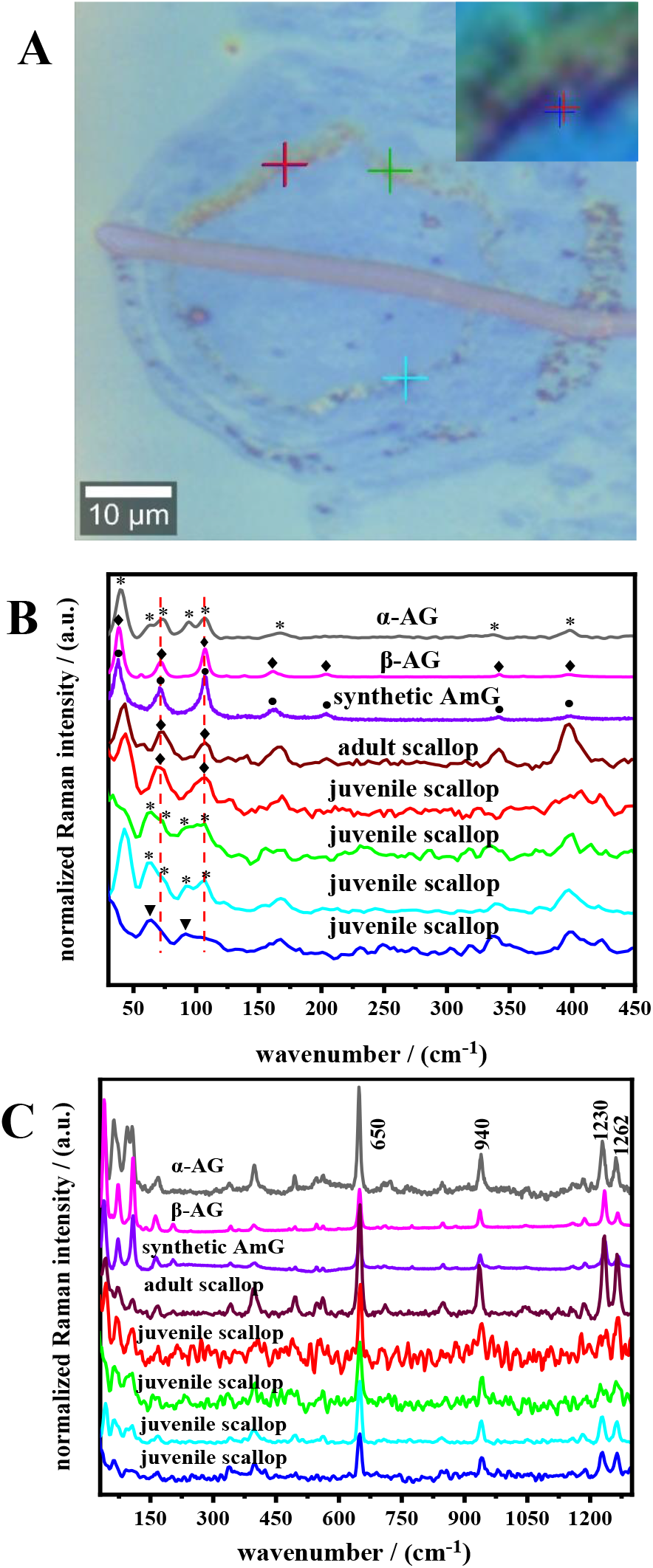
Micro-Raman analysis of the mirror region of the eyes of the juvenile scallops with size of 2.5 mm. (A) Optical microscopy image of the eyes of 2.5 mm scallops. (B) The low-wavenumber and (C) full wavenumber range Raman spectra of synthetic guanine references and the concave mirror region of scallop eyes. The Raman spectra of the synthetic α-guanine, β-guanine and amorphous guanine were identified by black, rosy red and purple curves respectively. Raman spectrum of the guanine in the concave mirror region of the eyes of adult scallops was marked with brownness curve, while the Raman spectra of the concave mirror region of the eye of 2.5 mm scallops marked with different cross markers were displayed by red, green, sky-blue and blue curves, respectively. The thickness of the slice samples of the biogenic samples was 200 nm, and the slices were placed on the carbon film of 100 mesh.

As shown in Figure 3, the characteristic peaks at 72 cm^-1^, and 107 cm^-1^ of β-AG were present at the position of the concave mirror region of the eyes of 2.5 mm juvenile scallops marked in red, which might assign to β-AG. In comparison, the characteristic peaks at 63 cm^-1^, 73 cm^-1^, 94 cm^-1^ and 107 cm^-1^ of α-AG appeared at the locations marked in green and sky blue of the concave mirror region of the eyes of 2.5 mm juvenile scallops, indicating the presence of α-AG. However, the peaks at 63 cm^-1^ were significantly higher than those at 73 cm^-1^ for the two Raman spectra at the locations marked in green and sky blue of juvenile scallops, very different from the Raman spectrum of standard α-AG. Interestingly, two obvious bands at 63 cm^-1^ and 94 cm^-1^ could be observed from the Raman spectrum of area marked in blue, while two other typical bands at 73 cm^-1^ and 107 cm^-1^ were broad or not obvious and a broad peak was present in between 94-116 cm^-1^. In comparison, the Raman spectra at different positions of the concave mirror region of the eyes of adult scallops showed pure β-AG **(Figure S3)**.

Thus we propose that there probably exists an unknown intermediate phase, biogenic amorphous phase of guanine and α-AG at these three locations, and α-AG is the minor phase at the location marked in blue considering the typical three of the four peaks of α-AG at short wavenumbers are not present. Considering that the synthetic AmG has short-range ordered structure similar to that of β-AG,^[30]^ it is reasonable that synthetic AmG and β-AG have similar Raman spectra. However, biogenic amorphous guanine shows different Raman spectra, typically, a strong vibration band at 63 cm^-1^ and broad bands in between 93 and 116 cm^-1^, different from synthetic AmG, α-AG and β-AG. In comparison, the concave mirror region of the eyes of adult scallops were fully developed and the amorphous guanine have crystallized into twining β-AG platelets. Therefore, no transformation from amorphous guanine to α-AG occurred during the sample preparation process for the adult scallop eyes. Thus, we suggest that the amorphous guanine is present in the concave mirror region of the juvenile scallops. And the amorphous guanine may function as an intermediate precursor in the eyes of juvenile scallops during the mineralization process.

### 2.3. Twinning crystals in the eyes of juvenile scallop (3 mm)

The eyes of 3 mm-sized scallops were also characterized to track the formation of twinning guanine platelets. Scallop eyes at this stage were well developed. The thickness of the concave mirror was about 8 μm, shown as width in the TEM image of the semi-thin transverse cross section **(Figure 4**A_1_), which was much wider than the width (2.6 μm) of the concave mirror region of 2.5 mm scallop (Figure 3A_1_). Many crystals still remained in the mirror region (Figure 4), although a large number of crystals were lost at both ends of the concave mirror during the sectioning process. In addition, some crystal platelets split into several pieces (Figure 4A_3_), probably due to the cutting force of microtome. Inside the concave mirror, except for the preexisting granular retina, novel continuous strips of tissue marked with green arrows were also observed **(Figure S4)**. Combining with the ultrastructures of the concave mirrors of the adult scallop eyes reported in the previous literature, we deduce that these continuous strips of tissue are the proximal retina.^[12]^ In comparison, the TEM images of the concave mirror region of adult scallops showed transverse cross sections of twinning guanine platelets with higher density than the cytoplasm in circumference area and the white holes indicate the loss of crystals **(Figure S5)**.

**Figure 4.**
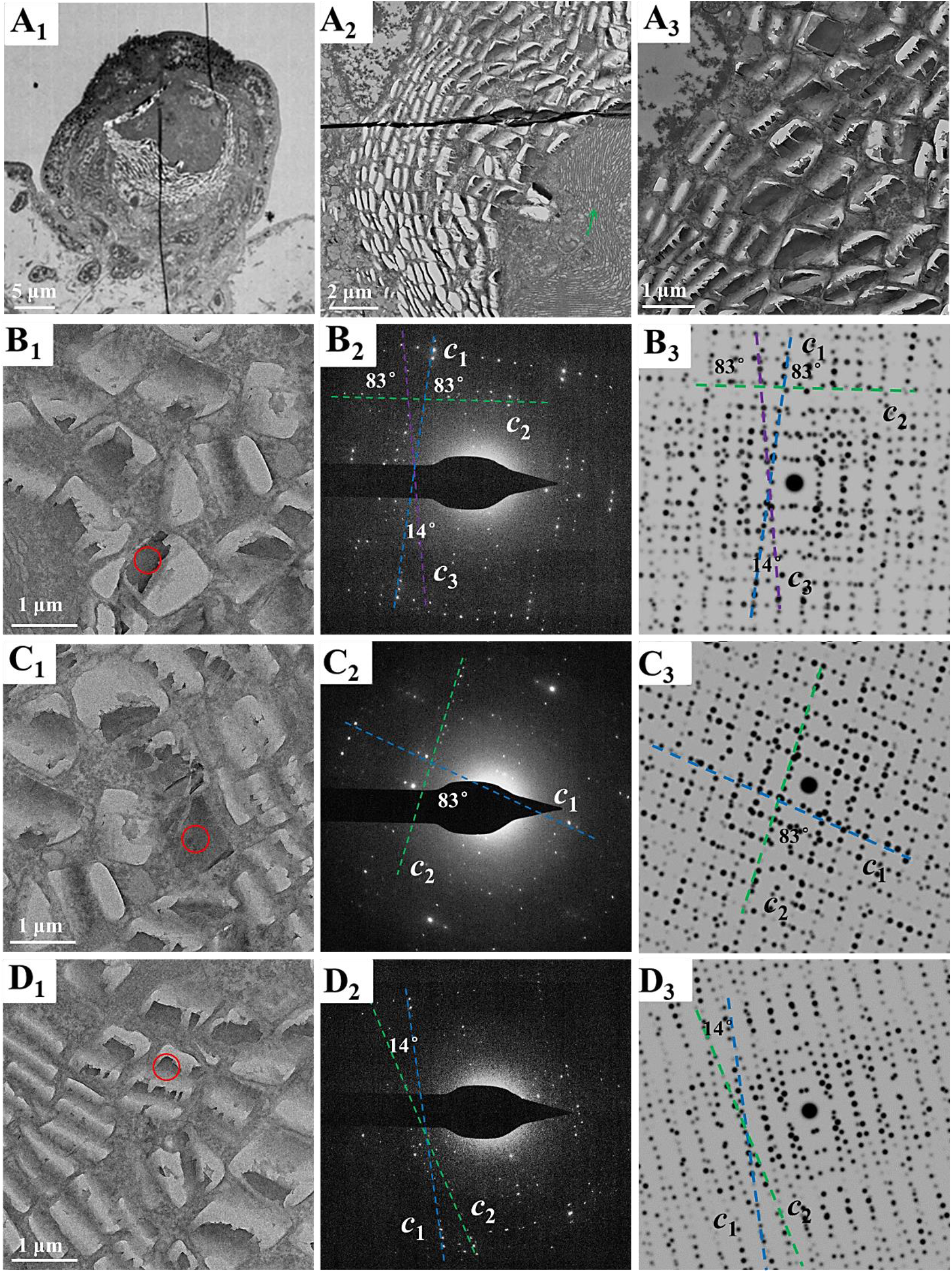
TEM images and SAED patterns of the guanine crystals in the eye of 3.0 mm juvenile scallops; the experimental SAED patterns (B_2_, C_2_, D_2_) and simulated SAED patterns (B_3_, C_3_, D_3_) of the guanine crystal in (B_1_, C_1_, D_1_). (A_1_) The eye with low magnification. (A_2_, A_3_) Zoomed-in of the region of concave mirror in (A_1_).

Diffraction spots along two or three different directions can be observed from each SAED pattern of the guanine crystal platelets in the eyes of 3 mm juvenile scallops (Figure 4). The d-spacing of the diffractions spots on each row was about 1.83 nm, indexed to the (001) plane of the β-AG. These results indicate that the platelets in the 3 mm scallop eyes are twinning β-AG crystals with two or three *c* axes, instead of single crystalline β-AG shown in Figure 3. The twinning angles of most crystals were 83° and 14°. As shown in the Figure 4B, the twinning guanine platelet had three *c* crystal axes. The twinning angles between *c*_1_ and *c*_2_ diffraction vectors, and *c*_2_ and *c*_3_ diffraction vectors were all 83°, while the twinning angle between *c*_1_ and *c*_3_ diffraction vectors was 14°, which were completely consistent with the twinning angles of twinning guanine crystals found in the eyes of adult scallops.^[18]^ For the marked guanine crystal in Figure 4C, the twinning angle between two *c* diffraction vectors was 83°. Furthermore, the guanine crystal shown in Figure 4D was a twinning crystal with twinning angle of about 14° between two *c*-axes. For all the characterized twinning crystals, the diffraction spots along *c*_1_ diffraction vector were generally stronger and denser than those along direction of *c*_2_ and *c*_3_ diffraction vectors. The crystallization degrees of guanine crystals along the directions of *c*_2_ and *c*_3_ axes were weaker than that along *c*_1_ diffraction vector. Considering that only single crystalline β-AG formed in the eyes of 2.5 mm-juvenile scallops, twinning β-AG crystals found in the eyes of 3.0 mm-juvenile scallops are supposed to be formed via single crystalline β-AG. We assume that a second crystal forms on the original single crystal nanoplatelet and twinning crystals start to form in the scallop eyes at this growth stage (3.0 mm). Simulated SAED patterns of twinning guanine microplatelets were shown in Figure 4, which were totally consistent with the experimental SAED patterns. The twinning angle (83° and 14°) of the guanine twinning microplatelets in the scallop eyes may be formed by the attachment of G-quartet (G4) to the (100) face, which induces the nucleation and growth of the second layer of guanine crystals.^[31]^ The stacked growth of crystal layers with different orientations probably induce the formation of twinning guanine crystals.

The twinning guanine microplatelets were composed of multiple stacked layers, rather than a single monolith **(Figure 5)**. At least four layers can be observed from the edges of the cubic microplatelet (Figure 5A_2_), which show a twinning crystal while the twinning angle between two *c*-axes was 83° according to the SAED pattern. Some twinning guanine platelets exposing side faces, transverse cross sections, show multilayers in Figure 5B and **Figure S6**. Generally, the platelets have no less than four observable layers. There is clear space between the layers, which were supposed to be formed during the microtome process. The transverse cross section of the guanine crystal in Figure 5B_2_ shows four superimposed layers. The thickness for the monolayer of the twinning guanine platelets was measured to be about 14±2 nm in average according to the TEM images shown in Figure 5 B and Figure S6 if we consider there are four layers. The total thickness of each twinning crystal is about 75±10 nm, which exceeded the sum of the thicknesses of the individual crystal layers because there is space between the layers. We assume that the measured crystal thickness of the twinned platelets of 3 mm scallops was bigger than the actual size due to the gaps caused by the cutting force. The thickness of monolayer is reliable since a lot of transverse cross sections of guanine microplatelets were applied for the measurements (Figure S6). According to the literature, the thickness of the twinning guanine microplatelets in the adult scallops is 75 nm.^[12, 26]^ The SAED pattern shows that the d-spacings were 1.82 nm and 0.32 nm in Figure 5B_2_, corresponding to (001) and (100) faces of β-AG, respectively. Furthermore, the wide and narrow sides are parallel to the (100) and (001) plane of β-AG, separately, which is consistent with the results of twinning β-AG microplatelets exposing large (100) face.^[18]^ Thus, the results for the twinning β-AG microplatelets formed in the junior scallops are consistent with the twinning β-AG platelets with three *c*-axes with twinning angles of 83° and 14° found in the adult scallops.^[18]^

**Figure 5.**
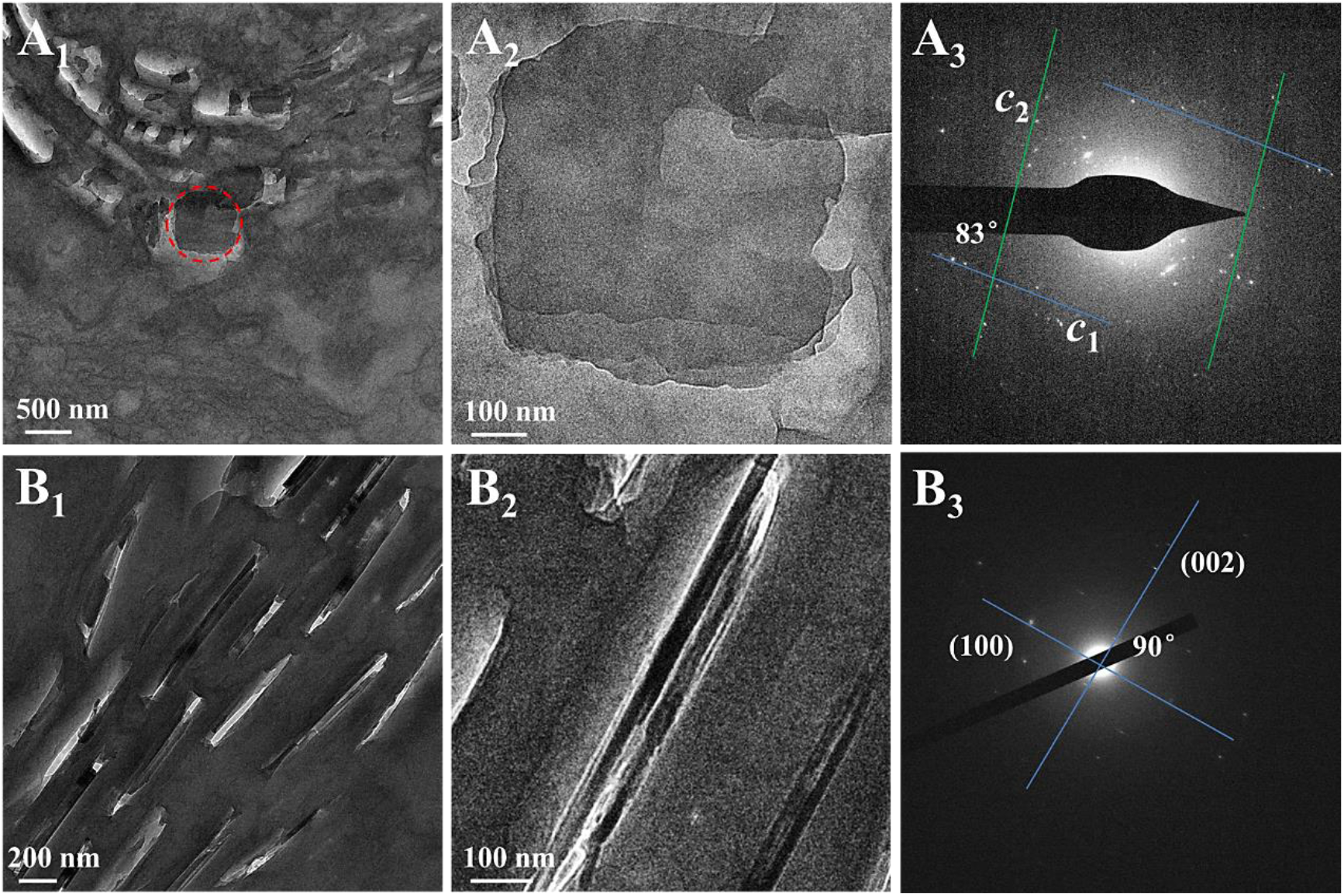
Cryo-TEM images and SAED patterns of twinning guanine microplatelets in the eyes of 3.0 mm juvenile scallop. (A_1_) the guanine twinning microplates exposing large crystal faces, (A_2_, A_3_) square mciroplatelet marked with red dotted circle in (A_1_) and SAED pattern, (B_1_, B_2_) the guanine crystals exposing side faces and the SAED pattern (B_3_) of the crystal shown in (B_2_). The crystal layers of guanine crystals shown in B_1_ and B_2_ were partially split during the cutting process of microtome. The thickness for the monolayer crystal of twinning guanine microplatelets was about 14 ± 2 nm.

## 3. Discussion

Wagner and collaborators proposed that guanine nucleates from a disordered precursor phase and the crystals grow by oriented attachment of partially ordered nanogranules into twinning platelets and then prismatic β-AG single crystals.^[19]^ As we all know, the square guanine microplatelets with ordered assembly in the adult scallop eyes are twinning β-AG crystals.^[18]^ The formation process and the crystalline states of twinning β-AG crystals in the eyes of juvenile *Yesso* scallops were investigated in detail in this work.

α-AG was found in the sections of juvenile scallop eyes according to Raman analysis in this work. We propose that α-AG may be formed via transformation from amorphous guanines during the sample preparation process, since α-AG is the thermodynamically most stable phase of anhydrous guanine. Micro-Raman spectroscopy in the concave mirror region of juvenile scallops detected not only the Raman signal of β-AG but also the characteristic peak of α-AG at several locations from the juvenile scallops with size of about 2.5 mm, which was supposed to be formed via transformation from amorphous guanine during the sample preparation process, providing indirect evidence for the presence of amorphous guanine precursors. Furthermore, we find a strong vibration band at 63 cm^-1^ and a very broad band in between 93 cm^-1^ and 116cm^-1^ from the Raman spectra of the juvenile scallops, which is a particular feature, different from the synthetic amorphous guanine (Syn. AmG), β-AG or α-AG. We propose that these novel features in the Raman spectra indicate the present of biogenic amorphous guanine (Bio. AmG). This is the first time to find direct proof for the presence of biological amorphous guanine in the biological organisms, even though undirected proof was shown from previous work.^[24]^ It was proposed that the β-AG microplatelets are transformed from amorphous guanine because guanine monohydrate was found from the fish scales,^[24]^ while guanine monohydrate was supposed not to be present in the fish scales.

TEM images indicate that single crystalline nanoplatelets of β-AG exposing (100) plane exist in the concave mirror region of the eyes of the juvenile scallops with size of about 2.5 mm. The existence of a large number of single crystalline β-AG nanoplatelets in the concave mirror region of the eyes of juvenile scallops with size of 2.5 mm, as the earliest crystalline stage, indicates that the twinning β-AG nanoplatelets formed at a later stage (3.0 mm juvenile scallops) and also in the adult scallops are formed via the single crystalline β-AG nanoplatelets.

Based on the results of TEM, SAED and Raman spectra, the crystallization mechanism of biogenic twinning guanine microplatelets in scallop eyes were proposed **(Scheme 1)**. Firstly, oval band region with layered framework form in the developing eyes in the very early stage, where the concave mirror composed of guanine crystals would form in the later stage. Secondly, guanine molecules would concentrate and amorphous guanine start to precipitate in the layered framework of the oval band region. Thirdly, a first layer of single crystalline β-AG nanoplatelets exposing (100) crystal face form via direct transformation via amorphous guanine, or, probably via dissolution and recrystallization process. G-quartet (G4) assembly may attach on the single crystalline β-AG nanoplatelets with a certain angle via interlayer π-π stacking, that is, the *b*-axis of β-AG nanoplatelets is parallel to z_1_ direction. A second nucleation proceed on the G4 and there forms a second nanoplatelet with *b* axis parallel to z_1_ or z_2_ direction, forming twinning crystals with twinning angle of 83°between the two *c* axes, or two-layered single crystalline nanoplatelets. Furthermore, a third thin nanoplate exposing (100) plane may form on the top of the second layer nanoplatelets with the *b* axis parallel to the z_1_ direction of G_4_. Thus, twinning β-AG nanoplatelets with three *c* axes with twinning angles of 83° between *c*_2_ and *c*_3_ diffraction vectors and 14° between *c*_1_ and *c*_3_ diffraction vectors were formed. The twinning β-AG microplatelets with twinning angle of 14° may be formed by the absence of *c*_2_ diffraction vectors. With a similar mode, multiple layers may form on the twinning β-AG nanoplatelets, and the nanoplatelets could grow to twinning β-AG microplatelets. Four or five layers were observed according to the TEM images of the twinning β-AG microplatelets, while the layer number of guanine crystal was bigger than the number of twinning angles. It is reasonable since a new layer crystal may have c axis the same as the original *c* axis or with a certain angle from *c*_1_ axis (83° or 14°). In the end, there are three types of twinning guanine in the concave mirror region in the later development stage of scallop eyes, twinning guanine microplatelets with two *c* axes with twinning angle of 83° or 14°, and twinning crystals with three *c* axes with twinning angles of 83° and 14°, respectively.

**Scheme 1.**
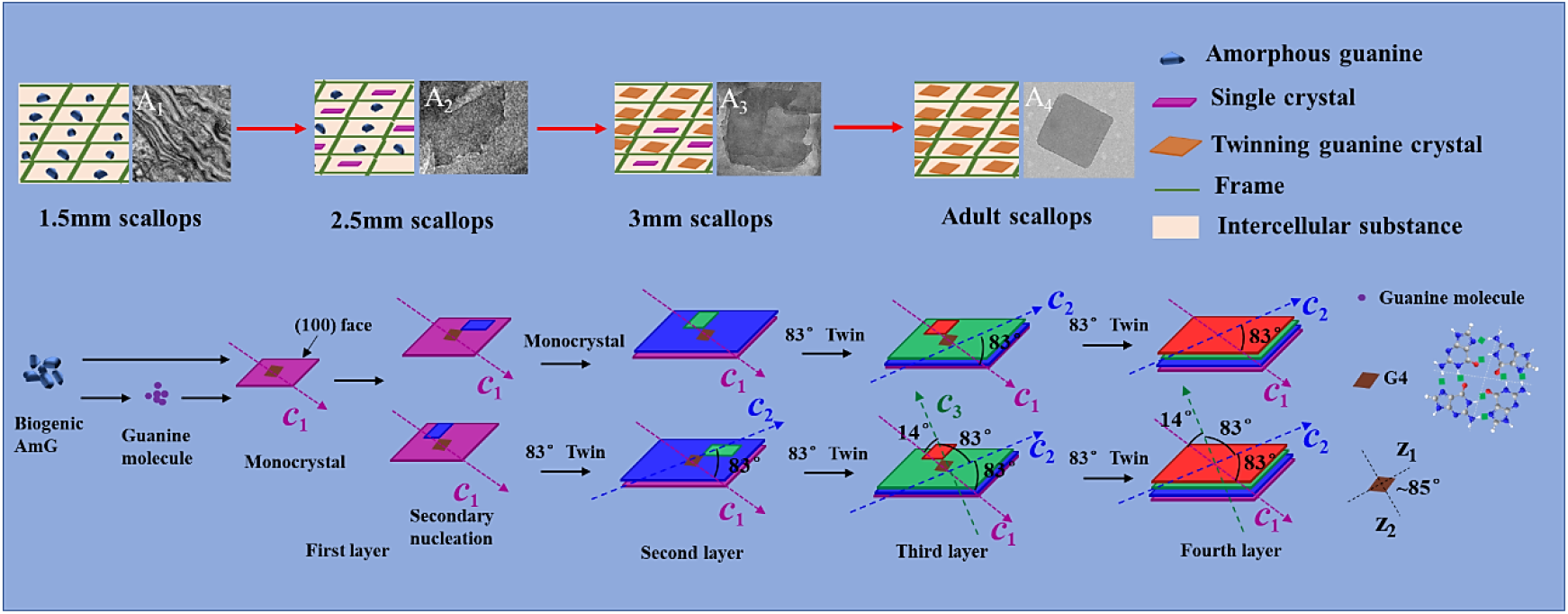
Schematic diagram of the formation process of the biogenic twinning microplatelets of the scallop eyes. The TEM image in A_4_ was reproduced with permission.^[18]^ Copyright 2017, Wiley-VCH.

## 4. Conclusion

The crystallization mechanism of twinning guanine platelets in the concave mirror region of the eyes of juvenile scallops were investigated in details in this work by using TEM and Raman spectroscopy. TEM indicates the transformation of guanine from single crystalline β-AG nanoplatelets to twinning β-AG nanoplatelets in the concave mirror region of the developing eyes of juvenile scallops with sizes from 2.5 mm to 3.0 mm. Besides β-AG and α-AG, a novel phase of guanine was found in the concave mirror region of the eyes of juvenile scallops with size of 2.5 mm according to the Raman spectra, which was proposed to be amorphous guanine, an intermediate phase as reservoir for guanine crystallization. Amorphous guanine may transform directly to thin layer crystalline β-AG nanoplatelets, or dissolve and recrystallize to β-AG nanoplatelets. α-AG was supposed to be transformed via amorphous guanine during the sample preparation process since α-AG does not exist in the eyes of adult scallops. A formation mechanism was proposed for the biogenic twinning β-AG microplatelets in the concave mirror region of scallop eyes based on the TEM and Raman spectra. Firstly, amorphous guanine is formed as a precursor phase in the layered frame-work cavity. Amorphous guanine would directly transform into crystalline β-AG, or, dissolve and recrystallize to single crystalline β-AG nanoplatelets. G4 assembly may attach onto the first layer monocrystalline β-AG nanoplatelets with a certain angle. Guided by G4 assembly, multilayer crystalline β-AG nanoplates nucleate and form twinning β-AG nanoplatelets, and then grow to microplatelets. And the *c* axes of the different layer crystalline β-AG nanoplatelets have certain angles from each other, 83º or 14º. Each layer of the β-AG nanoplatelets was calculated to be about 14 ± 2 nm. This is the first time to report the presence of amorphous guanine with direct proof and a clear formation mechanism of biogenic β-AG microplatelets is proposed. We assume that the formation mechanism of twinning guanine crystalline microplatelets with the same (100) plane, may represent a typical formation mechanism of twinning organic crystal plates sharing the same plane in the laboratories and organisms.

## 5. Materials and methods

### Materials

The glutaraldehyde solution (50%), sodium cacodylate trihydrate, uranyl acetate, sucrose and paraformaldehyde were bought from Sigma-Aldrich Co., LTD. Potassium ferrocyanide and uranyl acetate were supplied by Alfa Aesar. Ethyl alcohol was bought from Beijing hwrk chemical Co., Ltd. Spurr Embedding Kit was bought from Head (Beijing) Biotechnology Co., LTD. American cherry frozen embedding agent (OCT) was bought from Beijing solarbio science technology co., ltd. The *Yesso* scallops (*Patinopecten yessoensis*) were provided by the key Laboratory of Mariculture & Stock enhancement in North China sea, Dalian Ocean University. Juvenile scallops (with sizes from 1.5 mm to 3 mm) were collected from artificial nursery farm, and the adult scallops (7 cm in length) were collected from the artificial farm in the North China sea.

### Methods and characterizations

The eyes were dissected with a razor blade under microscope. The eyes of juvenile and adult *Yesso* scallops were fixed in a prepared fixative solution (4% paraformaldehyde, 2.5% glutaraldehyde and 2% sucrose in 0.1 M sodium cacodylate buffer at pH 7.4) at 4 °C for 24 h. The samples treated with the fixative solution were further fixed with a mixture of 2% osmic acid and 1.5% potassium ferrocyanide for 2 h, then were dehydrated in a series of gradient ethanol, and finally were embedded in Spurr resin.

The embedded samples were cut to semi-thin sections (200 nm) using a ultramicrotome (Leica EM UC7, Wetzlar, Germany), which were stained with uranyl acetate for later TEM analysis. The semi-thin sections of the eyes from juvenile and adult scallops were characterized by using TEM (FEI Talos F200X) and cryo-TEM (JEM-2100, equipped with Gatan 626-70 cryo-holder). Semi-thin sections of juvenile scallops were also tested by confocal Raman microscopy (alpha300R, WITec GmbH) using a diode-pumped solid-state laser (532 nm, cobalt Laser). Besides, after embedded in OCT and frozen with liquid nitrogen, serial 5 μm thick sections of adult scallop’s eye were obtained using a Leica CM1950 freezing microtome (Leica, Wetzlar, Germany) for Raman spectroscopy testing (alpha300R, WITec GmbH).

## Acknowledgements

This research was funded by the National Natural Science Foundation of China (Grant No. 21877009), and Beijing Institute of Technology Research Fund Program for Young Scholars. This work was also supported by the facilities at the Analysis & Testing Center, Beijing Institute of Technology.

## Author contributions

Conceptualization: D.G., Y.M. Methodology: D.G., Y.M., Y.L., X.W., C.F. Investigation: D.G., Y.L., X.H., H.Z. Characterization: D.G., Y.M., L.B., Y.L., X.H. Data analysis and processing: D.G., Y.M., X.H. Writing—original draft: D.G., Y.M. Writing—review & editing: D.G., Y.M.

## Notes

### Competing Interest Statement

The authors have declared no competing interest.

