## Supplementary Material for "Formation process of the twinning β-form anhydrous guanine platelets in the scallop eyes"

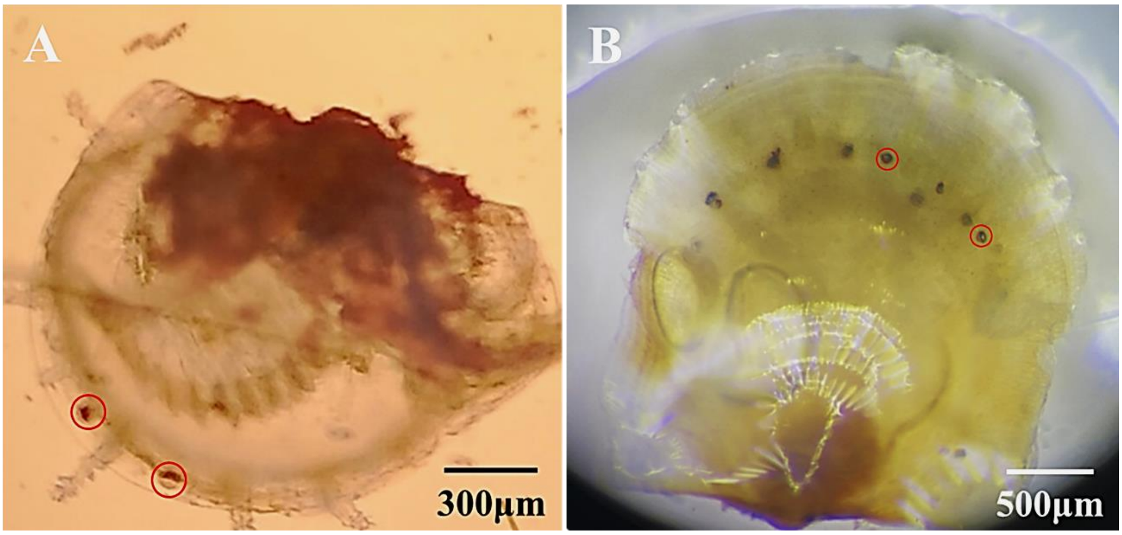


**Figure S1.** Photographs of juvenile scallops. (A) larvae scallops with size of 1.5 mm. (B) juvenile scallops with size of 2.5 mm. The eyes were marked with red circles. White reflections could be seen in the eyes of juvenile scallops with size of 2.5 mm.

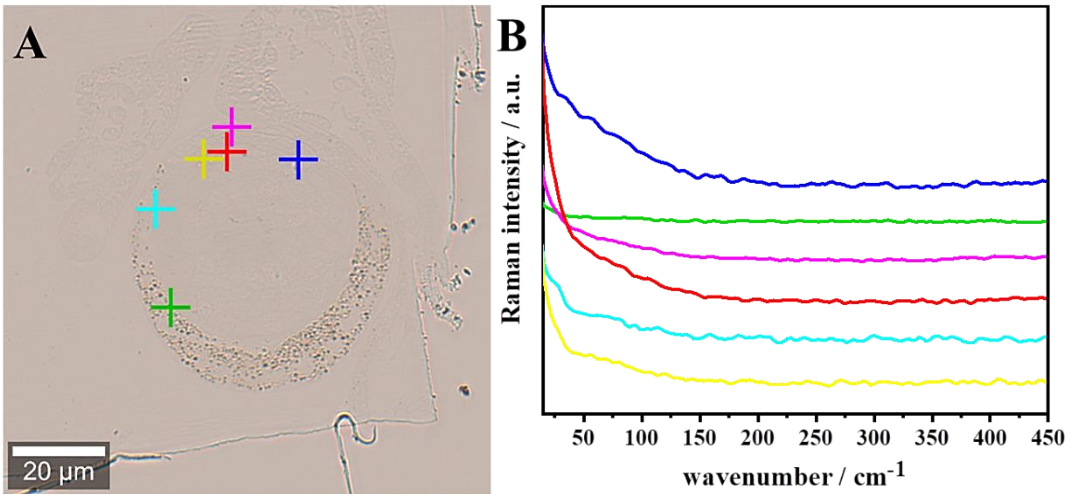


**Figure S2.** Optical microscopy image of the eyes of juvenile scallops with size of 1.5 mm (A) and corresponding low-wavenumber Raman spectra (B).

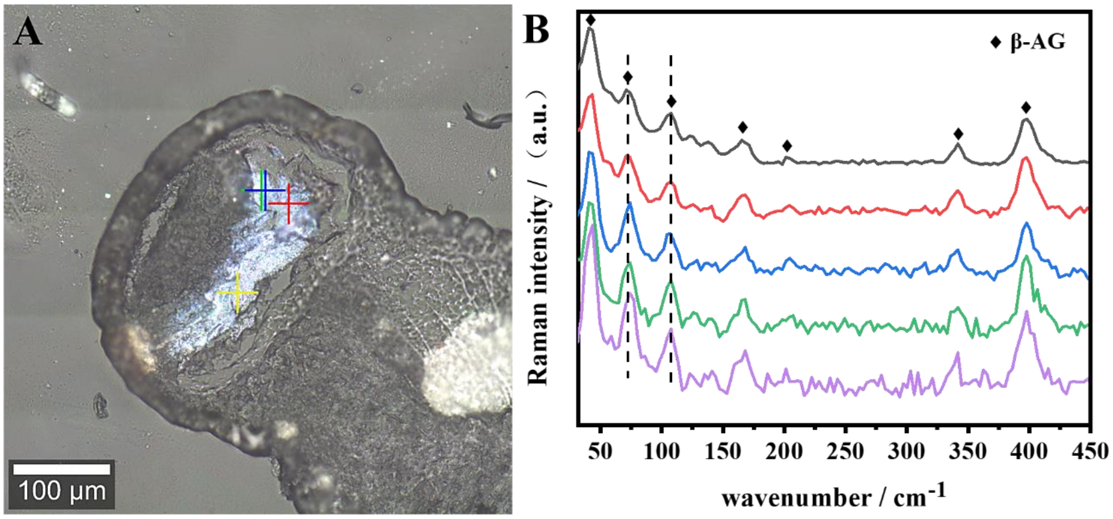


**Figure S3.** Optical microscopy image of the eyes of adult scallop **(A)** and corresponding low-wavenumber Raman spectra (B). All the characteristic peaks in the spectra are assigned to β-AG. The thickness of the slice samples of the biogenic samples was 5μm, and the slices were placed on the slide.

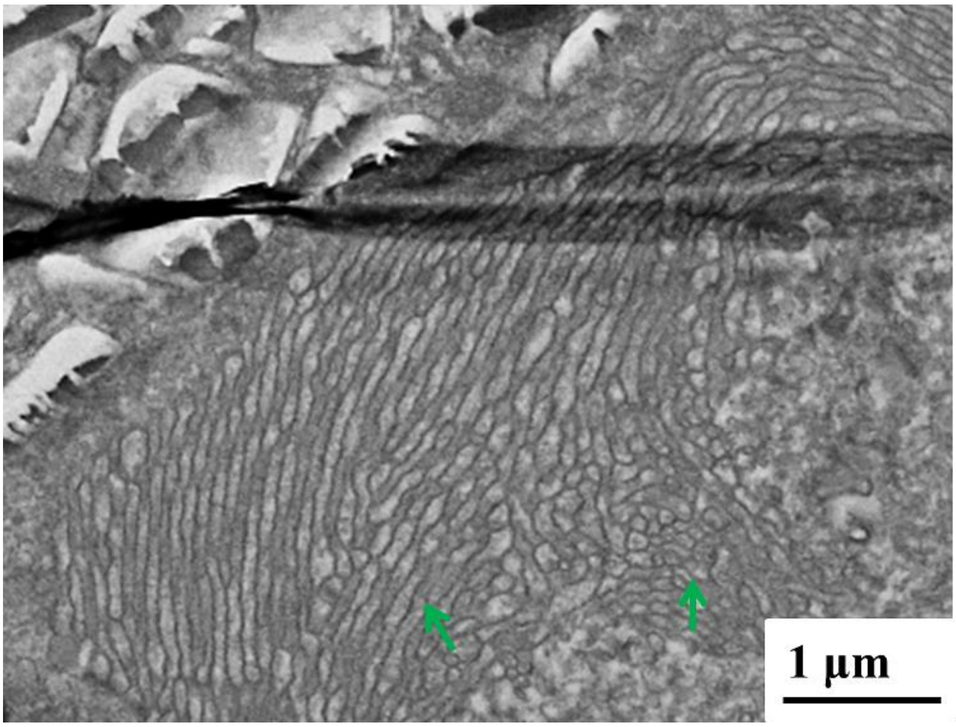


**Figure S4.** TEM image of the eye of juvenile scallop with size of about 3.0 mm. The continuous strips of tissue marked with green arrows are supposed to be the proximal retina.

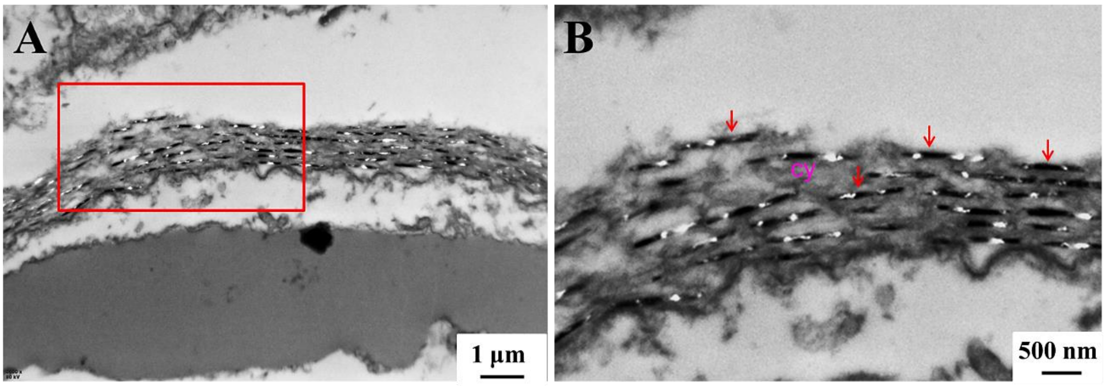


**Figure S5.** TEM images of guanine crystals in the concave mirror region of adult scallop eyes. (B) zoomed-in image of the region of concave mirror in red box of (A). *cy* cytoplasm.

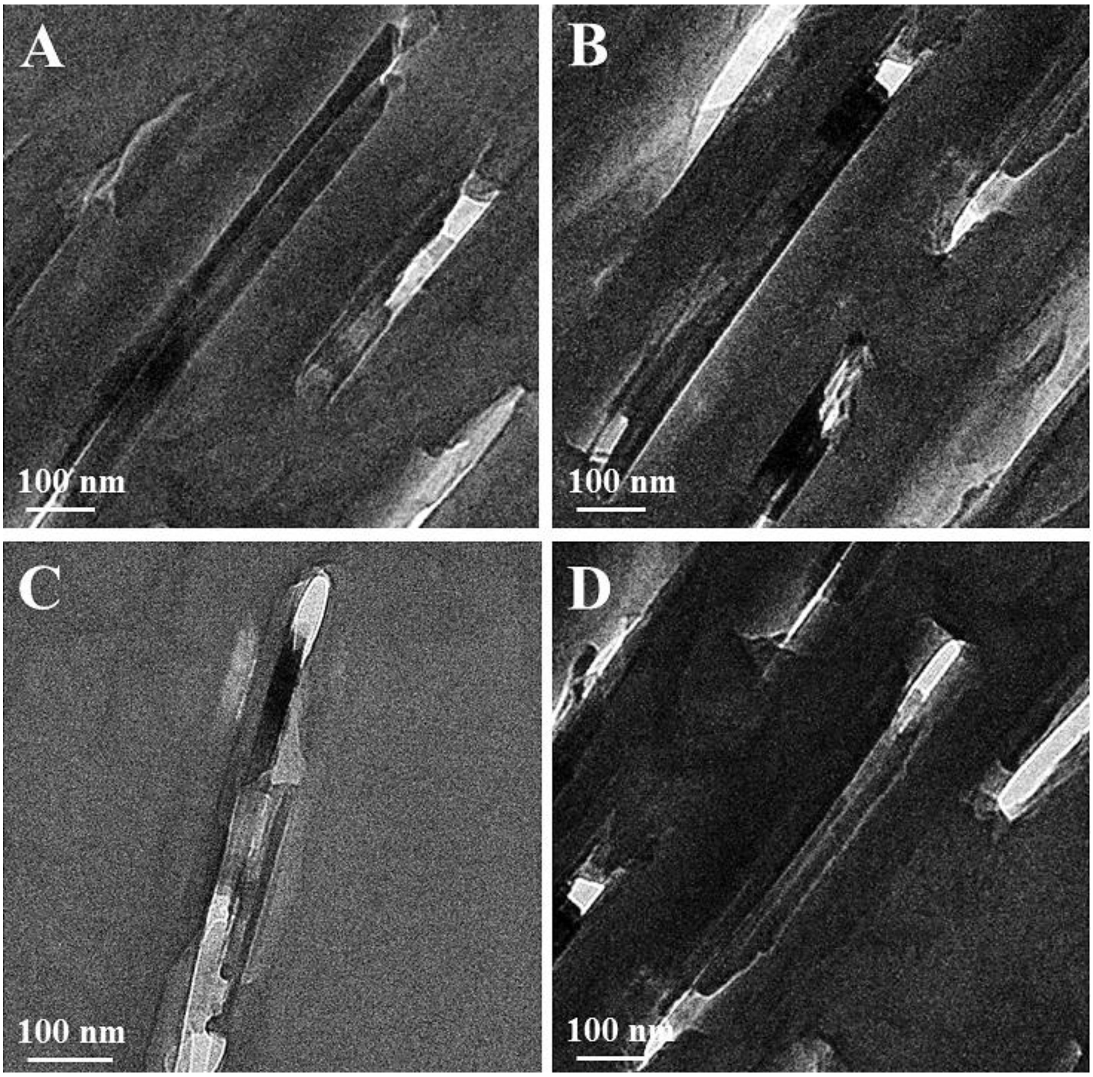


**Figure S6.** Cryo-TEM images of the guanine crystals exposing side faces.
